# Random folding drives the emergence of topologically associating domains in chromatin three-dimensional structure

**DOI:** 10.1101/2022.07.08.499343

**Authors:** Luming Meng, Qiong Luo

## Abstract

Domains are units of genome organization. Due to the seemingly irreconcilable difference between topologically associating domains (TADs) revealed by population-based biochemical studies and domains (sTADs) by single-cell imaging, finding a mechanism that simultaneously shape TADs and sTADs is challenging. Here we propose that TADs and sTADs are underlied by random folding of chromatin fiber heterogeneous in DNA density. On the hypothesis, we develop a model, termed RCHC, to yield chromatin structure ensemble from chromatin accessibility data. Calculated ensemble enables our hypothesis to be validated by population-based and single-cell experiments. Our simulation confirms the independence between domain and compartment structures in genome organization and shows that RCHC can predict the chromatin reorganization during differentiation. We mechanistically prove that genome is organized randomly with biases introduced by DNA-encoded information.

**One-Sentence Summary:** The TADs emergence is underlied by random folding of heterogeneous chromatin fiber carrying nucleosome occupancy information.

## Introduction

Mammalian chromatin is an intricately organized polymer in the cellular nucleus and its spatial organization directly influences gene regulation (*1-4*). Different experimental techniques for interrogating chromatin organization, such as high-throughput chromosome conformation capture (Hi-C) and fluorescence in situ hybridization (FISH), provide distinct aspects of chromatin organization (*5-7*). The lack of a view of integrating these aspects renders the mechanism that shape chromatin organization still unresolved.

Hi-C experiments performed on millions of cells investigate genome organization on the population-averaged level (*6*) and show that topologically associating domains (TADs) are structural units of genome organization within which chromatin preferentially interacts with itself (*8*). TADs are generally conserved across cell types and their boundaries are typically enriched at cohesin and CTCF binding sites (*5, 9*). Experiments of depleting cohesin or CTCF from cells show substantial disappearance of TADs in Hi-C maps (*10, 11*), indicating a defining role of cohesin and CTCF in the TADs formation (*12*). These observations provide the basis for the cohesin-CTCF mediated loop extrusion (LE) hypothesis, the now widely accepted mechanism for the formation of TADs (*13*). In this hypothesis, cohesin progressively extrude chromatin fiber until blocked by a pair of CTCF arranged in a convergent orientation, and then cohesin and CTCF cooperatively stabilize the extruded chromatin region as a spatially insulated domain (*14*). The LE hypothesis implicitly assumes that population-averaged TADs are physical structures present in single cells.

FISH experiments directly image chromatin structures in single cells and reveal that genomes in single cells are indeed partitioned into TAD-like domains with sharp boundaries (*15*). However, these single-cell TAD-like domains are not conserved among cells. Their boundaries show significant cell-to-cell variation, with a preference for residing at CTCF binding sites. The preference trivially explains the salient feature of population-averaged TADs (i.e. the enrichment of TAD boundaries at CTCF sites) and implies that TADs represent an emergent property rather than physical structures existing in individual cells. Therefore, single-cell TAD-like domains and population-averaged TADs are totally different in nature. For clarity, we refer to single-cell TAD-like domains as sTADs. To test the role of cohesin in the sTADs formation, Bintu *et al*. performed the FISH experiment on the cells after cohesin depletion, a treatment that abolishes TADs in the population Hi-C map (*15*). They found that sTADs persist after cohesin depletion and reappear after mitosis in the absence of cohesin, indicating that cohesin is likely not necessary for both the maintenance and the establishment of sTADs. It seems that cohesin does not play a vital role in the formation of sTADs. Hence, the cohesin-CTCF mediated LE hypothesis will run into overwhelming difficulty in explaining the sTADs formation.

We suggest that a plausible mechanism for the emergence or formation of TADs should be compatible with not only population-averaged Hi-C observations but also single-cell FISH observations. Here, we propose a new hypothesis rooted in the fundamental features of sTADs (*15*), including the stochastic variability in sTADs among single cells and the preference of sTAD boundaries for positioning at CTCF binding sites.

### RCHC model for the formation of sTADs and the emergence of TADs

Chromosomes can be considered as flexible polymer chains. In polymer physics, folding a long and flexible polymer chain within an extremely confined space (such as cell nucleus) will inherently lead to the result that local compaction occurs randomly and extensively along the polymer chain. We envision that when chromosomes are folded in an interphase cell nucleus, the locally compacted regions on chromatin fibers are sTADs observed by FISH imaging and the uncompacted or less compacted regions correspond to the boundaries of sTADs.

In the simple model, the inherently random nature of the distribution of locally compacted regions along chromatin fiber likely contributes to the observed cell-to-cell variability in sTADs (*16, 17*), but involves a difficulty in explaining the preference of sTAD boundaries for residing at CTCF binding sites. To solve the problem, we attempt to find out the distinct changes to chromatin fiber when CTCT binds to it. Studies on the property of CTCF binding sites show that CTCF binding can create arrays of closely spaced nucleosomes in neighboring chromatin regions, while the center of CTCF binding site corresponds to a nucleosome-depleted region (*18-21*). The observation implies that CTCF binding has a high correlation with the non-uniform nucleosome distribution along chromatin fiber. Therefore, it is necessary to include the information on nucleosome distribution in our model. To this end, we treat chromatin fiber as heterogeneous polymer chain along which the nucleosome distribution is non-uniform. Because the nucleosome distribution is mirror of the distribution of DNA density (DNA sequence length per unit physic length or volume), here, we consider that chromatin fiber is heterogeneous in terms of DNA density. Finally, we propose that Random local Compaction of Heterogeneous Chromatin fiber (RCHC) gives rise to the formation of sTADs and the emergence of TADs (Fig. 1A).

**Fig. 1.**
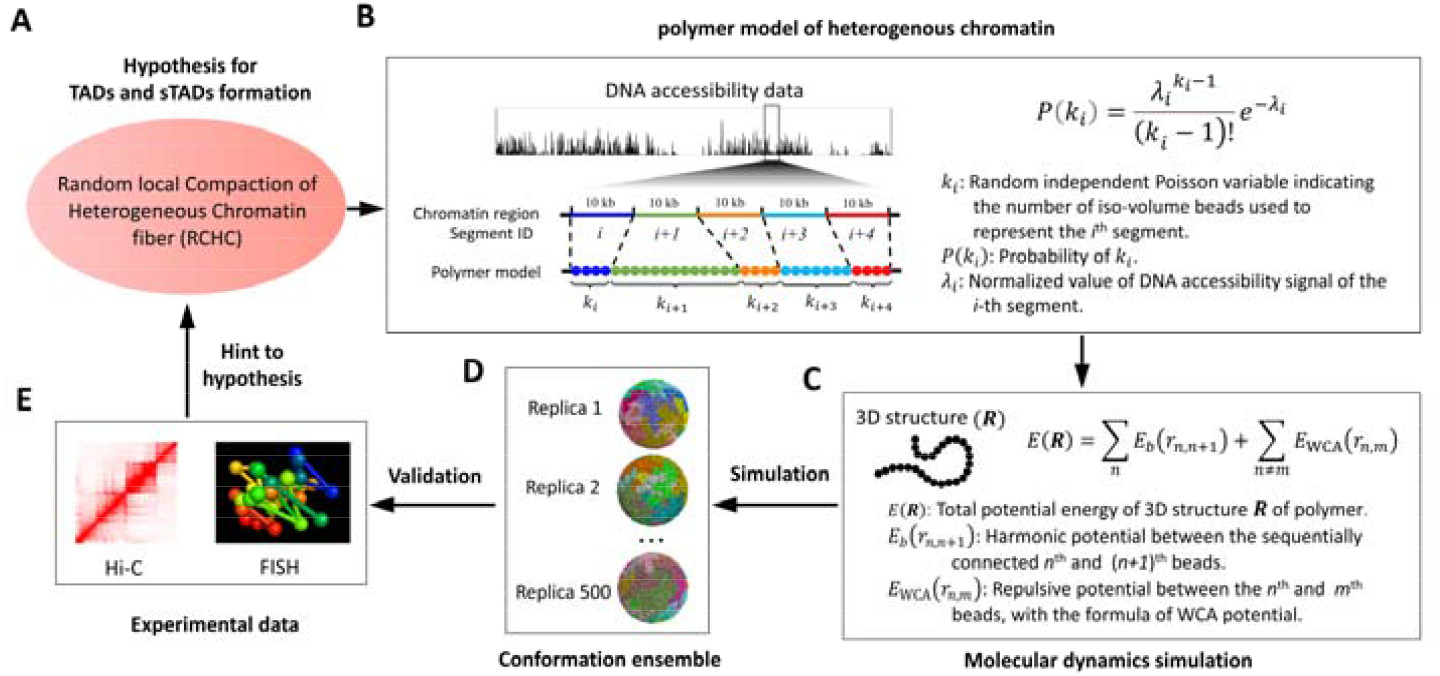
The RCHC model allows de novo prediction of an ensemble of 3D chromatin structure with chromatin accessibility data as input. (**A**) The hypothesis behind the RCHC model for the formation of sTADs and TADs. Notably, chromatin fiber is heterogeneous in term of DNA density. (**B**) A scheme of the approach for building polymer model of heterogeneous chromatin fiber. The chromatin region of interest is partitioned into consecutive 10-kb segments. The number *k*_*i*_ of coarse-grained beads used to represent the *i*^*th*^ segment is modeled as a random independent Poisson variable that is associated with chromatin accessibility of the *i*^*th*^ segment. Notably, beads are equal in physical volume but different in DNA sequence length. (**C** to **E**) Molecular dynamics simulation on random folding of polymer in a confined space (C) to produce a conformation ensemble consisting of 500 replicas for the chromatin region of interest (D). The conformation ensemble produced in (D) is compared to single-cell FISH data and population-averaged Hi-C data (E). Further details of the polymer model and MD calculations are given in the supplementary materials [(*24*), sections 1 to 2].

To test the RCHC hypothesis, we model heterogeneous chromatin fibers with self-avoiding polymer chains consisting of contiguous, coarse-grained beads (Fig. 1B). Notably, beads are same in physical volume but different in DNA sequence length. The central problem of building the polymer model of heterogeneous chromatin is how to quantify the DNA density (i.e. the represented DNA sequence length) of each equal-volume bead. To solve the problem, we turn it into another problem of determining the number *k* of equal-volume beads used to represent a given chromatin segment of fixed sequence length. We model *k* as a random independent Poisson variable which is associated with chromatin accessibility of the given chromatin segment.

The work flow of establishing polymer model of heterogeneous chromatin fiber is described in Fig. 1B. Specifically, we partition a chromatin region of interest into consecutive *N* 10-kb segments and assign a Poisson variable *k* to each segment. Then, we obtain a set of Poisson variable *K* (*k*_*1*_, *k*_*2*_,…*k*_*N*_) to build a polymer model for the chromatin region of interest. A certain *K* (*k*_*1*_, *k*_*2*_,…*k*_*N*_) corresponds to a certain DNA density distribution along the chromatin region of interest. To model the fact that the nucleosome distribution (i.e. the DNA density distribution) along chromatin shows cell-to-cell variation, we randomly generate a group of *K* (*k*_*1*_, *k*_*2*_,…*k*_*N*_) consisting of 10 members which represent ten different DNA density distributions of the same region. In other words, we build ten distinct polymer models for the same chromatin region of interest. For each polymer model, its stochastic folding in a confined space is simulated by molecular dynamics (MD) calculation (Fig. 1C) for times and starting from different random initial structures. Finally, we collect a conformation ensemble for the chromatin region of interest (Fig. 1D). The ensemble allows us to validate RCHC hypothesis with data from single-cell FISH and population-averaged Hi-C experiments (Fig. 1E).

Briefly, we propose a testable model, termed RCHC, for the formations of sTADs and TADs, which can de novo predict an ensemble of 3D chromatin structures with population-averaged chromatin accessibility data as input. There are two reasons that motivate us to use population-averaged chromatin accessibility data as the input data. First, multiple evidences show that the chromatin segments exhibiting high accessibility signal have relative low density of DNA and vice versa, revealing a high correlation between chromatin accessibility and DNA density (*22, 23*). Second, DNase-seq and ATAC-seq experiments have directly yielded a terrific amount of chromatin accessibility data for various cell types at the population level (*22, 23*). These reasons convince us to believe the promise that the RCHC model can predict chromatin organizations of various cell types under the laws of physics without priori assumptions.

### RCHC quantitatively predicts TADs at genome-wide scale

We apply RCHC to generate 3D structures of the chromatin region including 22 autosomes and an X-chromosome of K562 human erythroleukemia cell type and collect an ensemble containing 500 conformations, with the ATAC-seq data (*24*) from the Gene expression Omnibus (GEO) as input. Because the ensemble describes the random folding of heterogeneous chromatin, we denote it as hetero-ensemble(ATAC). We select a 5-Mb chromatin region (Chr5:109Mb-114Mb) as example for the quantitative comparison between the simulation and the population-averaged Hi-C data of K562 (*24*).

For each conformation, we measure contacts between each pair of 10-kb chromatin segments throughout the 5-Mb region and construct an individual contact matrix at 10-kb resolution (Fig. 2A). We average the individual contact matrices across the whole ensemble and obtain the ensemble-averaged contact matrix (Fig. 2C) which can be compared to the 10-kb resolution Hi-C contact-frequency matrix of K562 (Fig. 2B). Quantitatively, we identify domain boundaries from all contact matrices using separation score (*24*) which quantify the degree of spatial separation between upstream and downstream chromatin of a position (Fig.2). Notably, TAD boundaries that are visual evident in the Hi-C matrix always correspond to separation score peaks (Fig. 2B), supporting the validity of separation score for defining domain boundaries. Comparison between the Hi-C matrix (Fig. 2B) and the ensemble-averaged matrix (Fig. 2C) shows that the simulation can predict most TAD boundaries in the Hi-C matrix, indicating the validity of the RCHC model.

**Fig. 2.**
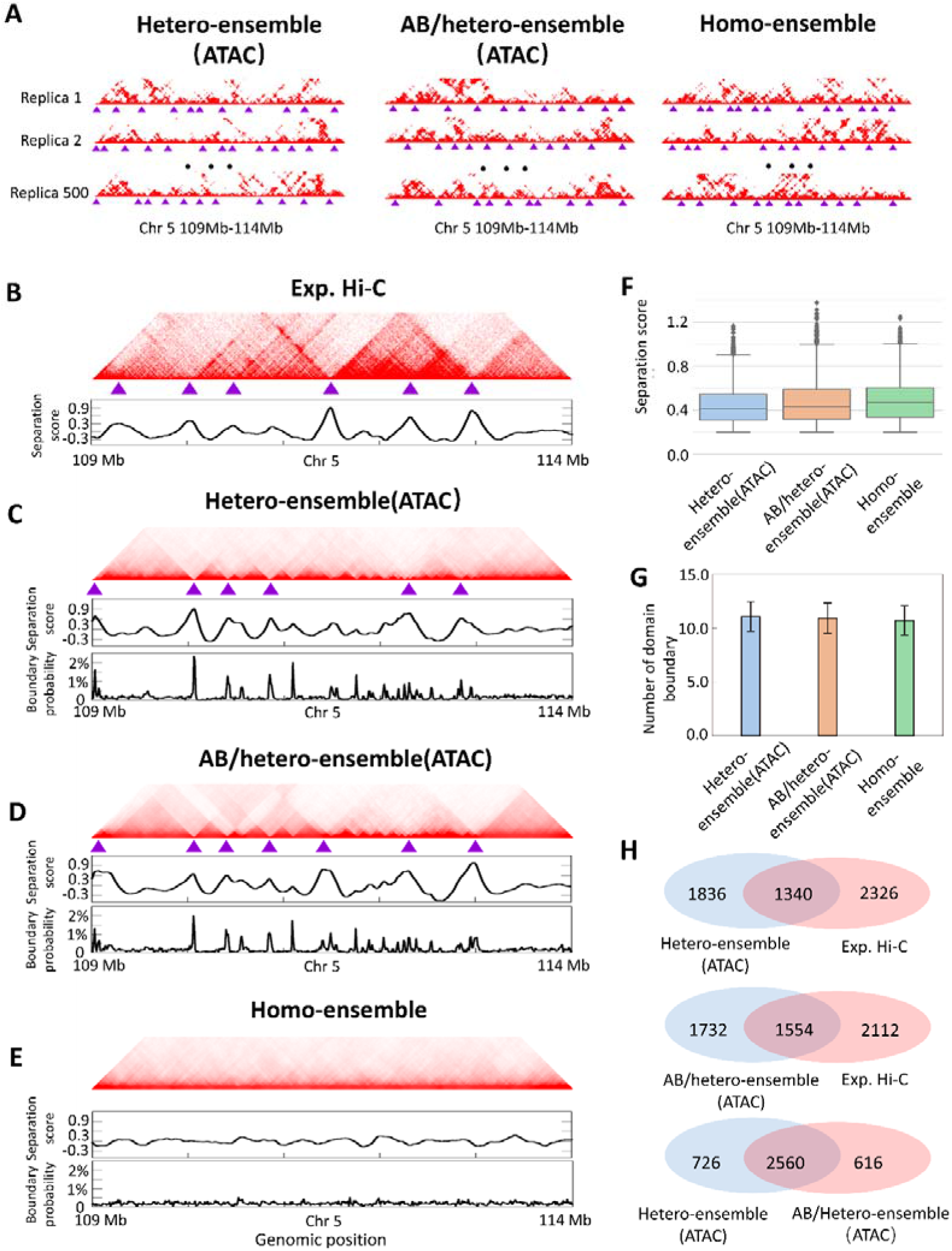
Validation of the RCHC model with Hi-C data. (**A**) Individual contact matrices of the 5-Mb chromatin region (Chr5:109Mb-114Mb) in K562 at 10-kb resolution, calculated from individual conformations within the three simulated ensembles: hetero-ensemble(ATAC), describing random folding of chromatin fiber heterogeneous in DNA density using ATAC-seq data as input; AB/hetero-ensemble(ATAC), describing the same random folding process as hetero-ensemble(ATAC) but under the interaction for A/B compartment formation; homo-ensemble, describing random folding of homogeneous chromatin fiber. (**B**) Top: Population-averaged Hi-C contact matrix for the 5-Mb chromatin region at 10-kb resolution [data from ref(*24*)]. Bottom: Separation sore for each genomic position in the Hi-C matrix. (**C** to **E**) Top: The 10-kb resolution ensemble-averaged contact matrices of the 5-Mb chromatin region for hetero-ensemble(ATAC) (C), AB/hetero-ensemble(ATAC) (D), and homo-ensemble (E). Each ensemble-averaged matrix is calculated by averaging individual contact matrices shown in (A) across the whole corresponding ensemble. Middle: Separation sore for each genomic position in the ensemble-averaged contact matrix. Bottom: Probability (fraction of the 500 individual conformations) for each genomic position to appear as a domain boundary. Purple triangles in (A-E) denote positions of domain boundaries identified by separation score. (**F**) Boxplots showing separation scores of domain boundaries in the individual contact matrices of the 5-Mb region. (**G**) The number of domain boundaries identified from each individual contact matrix of the 5-Mb region. (**H**) Overlap in domain boundaries throughout 22 autosomes and X-chromosome of K562 among the population-averaged Hi-C contact matrix, the ensemble-averaged contact matrices of Hetero-ensemble(ATAC) and AB/hetero-ensemble(ATAC). Further details of calculating individual and ensemble-averaged contact matrices, separation score, and overlap are given in the supplementary materials [(*24*), sections 3 and 5].

Furthermore, we examine the ensemble-averaged matrix of the intact chromatin region containing 23 chromosomes of K562 and find that the total number of domain boundaries is 3176 among which 42.2% share their genomic positions with TAD boundaries in the Hi-C map of K562 (Fig. 2H). However, the percentage decreases to 19.0 % when we randomly select 3176 genomic positions throughout the intact region consisting of 23 chromosomes of K562. These results show that the RCHC model can quantitatively predict TADs revealed by population Hi-C data, at genome-wide scale.

### RCHC reproduces single-cell FISH data

In principle, conformations in the hetero-ensemble(ATAC) can be considered as snapshots of chromatin organization in single cells. Hence, the measurements of individual conformations can be compared to the information obtained from single-cell FISH images. A glance of the individual contact matrices derived from conformations shows widespread existence of domain structures in each conformation and substantial variation in the positions of domain boundaries between conformations (Fig. 2A). Moreover, the measurement of the probability of each genomic position for appearing as a domain boundary across the 500 individual matrices shows that more than 98% of positions throughout the 5-Mb region (Chr5:109Mb-114Mb) display a nonzero probability (Fig. 2C). Notably, the curve of probability exhibits peaks at the positions aligning with TAD boundaries in the Hi-C matrix, indicating a preference of such positions for appearing as a domain boundary in conformations (Fig. 2B and 2C). These results demonstrate that the structural features of simulated individual conformations are exactly consistent with the features of single-cell chromatin organization revealed by FISH experiments (*15*).

Furthermore, we quantitatively assess the validity of the RCHC model using concrete FISH data. For each conformation in the hetero-ensemble (ATAC), we construct a distance matrix (Fig. 3A) by measuring spatial distances between each pair of 30-kb segments throughout the 2-Mb genomic region (Chr21:29.38Mb-31.33Mb) and average them to obtain the ensemble-average distance matrix (Fig. 3B). Notably, the simulated ensemble-averaged distance matrix (Fig. 3B) shows a high correlation with the average distance matrix derived from the FISH experiment (Fig. 3C)(*15*), with Pearson correlation coefficient of 0.935 (Fig. 3F). Moreover, domain structures identified from the simulated individual distance matrices adopt spatially segregated globular configurations (Fig. 3A), consistent with the appearance of sTADs observed in the super-resolution FISH experiment (*15*). These results provide a further evidence for the validity of the RCHC model.

**Fig. 3.**
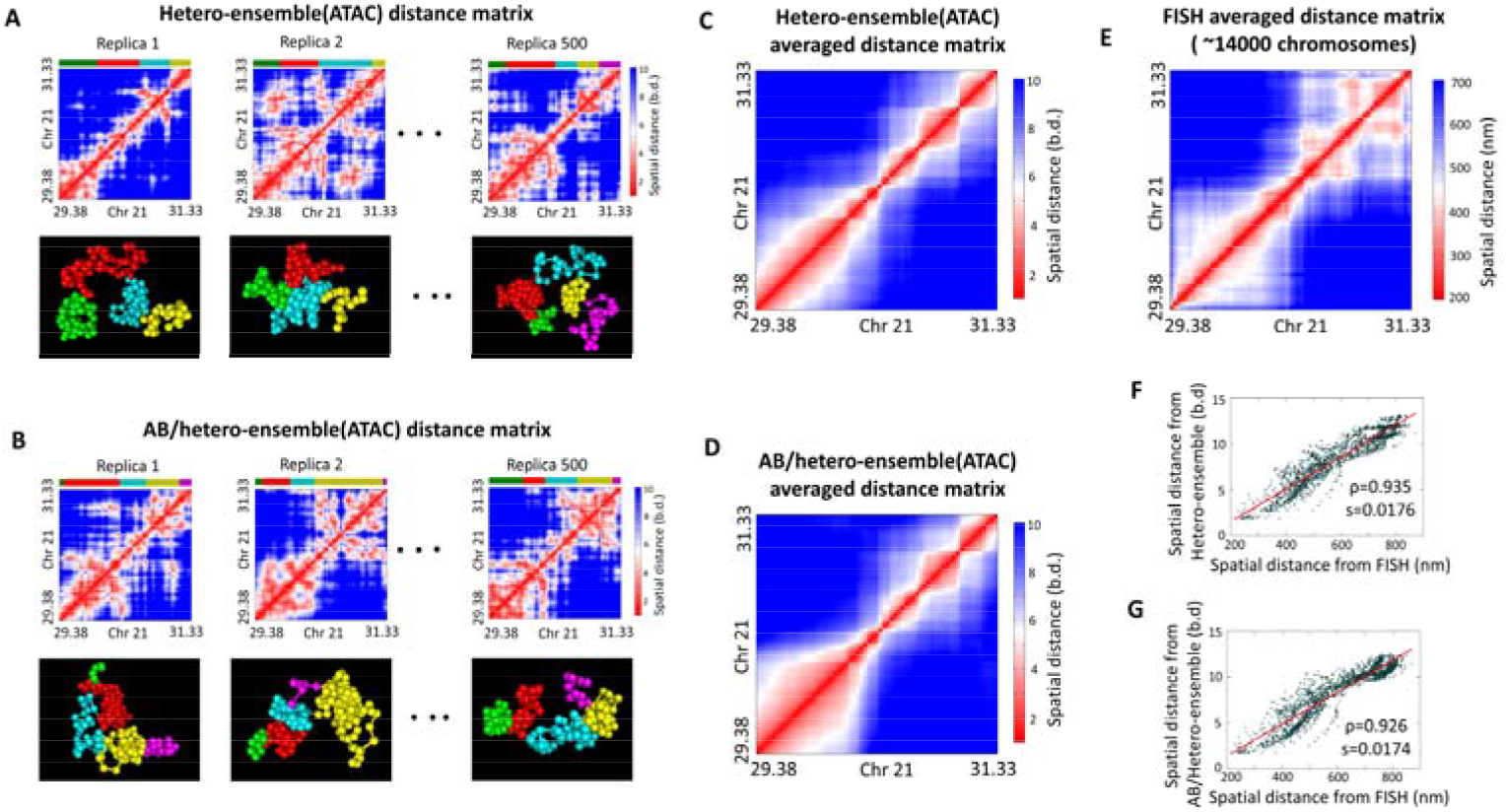
Validation of RCHC model with FISH data. (**A** and **B**) Top: Individual distance matrix of the 2-Mb genomic region (Chr21:29.38Mb-31.33Mb) of K562 at 30-kb resolution, calculated from individual conformations within hetero-ensemble(ATAC) (A) and AB/hetero-ensemble(ATAC) (B). Bottom: 3D configurations of the 2-Mb genomic region. The genomic regions marked in green, red, cyan, yellow, and purple correspond to domains identified by separation score. (**C** and **D**) The 30-kb resolution ensemble-averaged distance matrices of the same region for hetero-ensemble(ATAC) (C) and AB/hetero-ensemble(ATAC) (D), calculated by averaging the individual distance matrices shown in (A) and (B), respectively. (**E**) The experimental averaged distance matrix at 30-kb resolution across ∼14,000 FISH images of the same region [data from ref. (*15*)]. (**F**) Correlation between the distance matrices shown in (C) and (E). (**G**) Correlation between the distance matrices shown in (D) and (E). Further details of distance matrices calculation are given in the supplementary materials [(*24*), sections 4].

### Assessment of the practicability of the RCHC model

We assess the practicability of the RCHC model from three dimensions: scale of chromatin organization, input data source, and cell type. First, the above mentioned results of the hetero-ensemble(ATAC) prove that RCHC allows us to predict chromatin structure of high resolution at multiple scales: from several megabases to intact genome. Second, we investigate whether the sources of chromatin accessibility data (i.e. ATAC-seq or DNase-seq experiments) influence the performance of the RCHC model. We repeat the process of generating the hetero-ensemble(ATAC) but with the DNase-seq data of K562(*24*) rather than ATAC-seq data as input to produce the hetero-ensemble(DNase). The analysis of the hetero-ensemble(DNase) (Fig. S4 and S5) reaches the same conclusions of the hetero-ensemble(ATAC), indicating that the RCHC model is compatible for the two types of chromatin accessibility data from ATAC-seq and DNase-seq experiments, respectively. Third, we focus on whether the RCHC model is applicable to other cell types except for K562 erythroleukemia cell type. We apply RCHC to generate 3D structures of the chromatin region including 22 autosomes and an X-chromosome of IMR90 lung fibroblasts cell type with the ATAC-seq data(*24*) from the GEO as input and also collect an ensemble consisting of 500 conformations. Multiple comparisons between the RCHC simulated results and the experimental data of IMR90(*24*) again show that the RCHC simulation can predict the experimental observations of chromatin structure from population-averaged Hi-C and single-cell FISH experiments (Fig. S6 to S8). These results indicate that the RCHC model is practical in the field of genome biology.

### RCHC confirms the independence between local insulation and genomic compartmentalization in genome organization

Mammalian genome organization shows two fundamental features in interphase: genomic compartmentalization of chromatin into A/B compartments and local insulation of chromatin into domains(*13, 14*). An interesting question is: what the interplay is between local insulation and genomic compartmentalization in genome organization. Previous studies show that self-interaction between heterochromatic regions is the mechanism responsible for compartmentalization(*25*). We incorporate such interaction into the RCHC modeling of random folding of heterogeneous chromatin and generate a new conformation ensemble for K562, denoted as the AB/hetero-ensemble(ATAC). We perform a comparison between the AB/hetero-ensemble(ATAC) and hetero-ensemble(ATAC) to investigate the relationship between local insulation and genomic compartmentalization (Fig. 2).

We first find that the qualitative features of individual contact matrices, including the widespread existence of domains in individual conformations and high variation in domain boundaries among conformations, persist between the two ensembles (Fig. 2A). Quantitatively, the number of domain boundaries within individual conformations and boundary strength characterized by separation score also remain similar between the two ensembles (Fig. 2F and 2G). Next, we observe that the ensemble-averaged contact matrix of the 5-Mb region (Chr5:109Mb-114Mb) of the AB/hetero-ensemble(ATAC) also predicts most TAD boundaries in the Hi-C matrix of the same region (Fig. 2B and 2D). We further identify domain boundaries from the average contact matrices of the two ensembles throughout the whole region including 23 chromosomes. We find that the interaction for compartmentalization leads to a very slight increase in the total number of boundaries (from 3176 to 3286), with the number of pairs of overlapping boundaries between the two ensemble being up to 2560 (Fig. 2H). Among the total 3286 boundaries in the average contact matrix of the AB/hetero-ensemble(ATAC), 47.3% are present at the genomic positions aligning with the TAD boundaries of the Hi-C map, and the percentage is slightly higher than that of the hetero-ensemble(ATAC) (i.e. 42.2%) (Fig. 2H). Meanwhile, we compare the ensemble-averaged distance matrices of the 2-Mb genomic region (Chr21: 29.38-31.33□Mb) between the two ensembles and observe no notable difference (Fig. 3). Together, these results indicate that the interaction associated with genomic compartmentalization do not meaningfully alter local insulation of chromatin into domains.

Furthermore, we derive three Pearson correlation matrices of the 20-Mb region (Chr2:30Mb-50Mb) for K562, from the Hi-C matrix and the ensemble-averaged contact matrices of the AB/hetero-ensemble(ATAC) and the hetero-ensemble(ATAC), respectively (Fig. 4). Notably, the plaid patterns in the Hi-C correlation matrix corresponding to signatures of compartments are unambiguously predicted in the correlation matrix of the AB/hetero-ensemble(ATAC), while no plaid pattern can be observed in the correlation matrix of the hetero-ensemble(ATAC). These results demonstrate that self-interaction among heterochromatic regions indeed shapes compartments in genome organization.

**Fig. 4.**
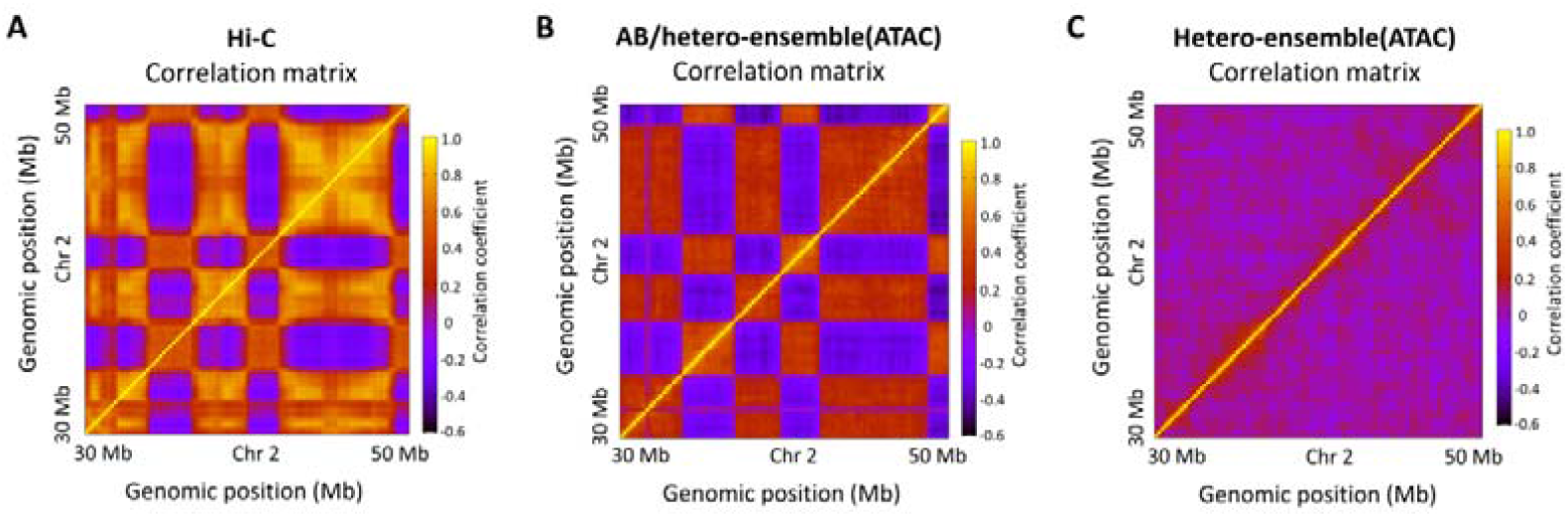
Compartment structures appear when the interaction for the compartment formation is taken into account in the simulation on random folding of heterogeneous chromatin fiber. (**A**) Pearson correlation matrix of the 20-Mb region (Chr2:30Mb-50Mb) of K562 derived from the Hi-C contact-frequency map [data from ref.(*24*)]. The plaid patterns are signatures of compartments. (**B** and **C**) Pearson correlation matrices derived from ensemble-averaged contact matrices of the AB/Hetero-ensemble (B) and Hetero-ensemble (C). Further details of calculating Pearson correlation matrix are given in the supplementary materials [(*24*), sections 6].

Overall, our simulations show that compartments and domains in genome organization are formed by independent mechanisms, which coincide with the observation that compartmental alteration do not correlate with TAD changes during differentiation process(*26*). Notably, the RCHC model with the inclusion of the interaction for compartmentalization shows an improved prediction of genome organization. The same conclusion can be obtained from the comparison of other two simulated ensembles which are generated by using the DNase-seq data as input (i.e. the AB/hetero-ensemble(DNase) and the hetero-ensemble(DNase) (Fig. S4 and S5).

### RCHC simulation reveals that genome organization is inherently random at submegabase scale

The simulations on random folding of heterogeneous chromatin fiber with and without the interaction for compartmentalization indeed show the formation of sTADs and the emergence of TADs. We next investigate the question: what is the consequence of randomly folding homogenous chromatin with or without the interaction of compartmentalization. By setting all the Poisson variables in *K* (*k*_*1*_, *k*_*2*_,…*k*_*N*_) equal to 1 in the RCHC model (Fig. 1B), we obtain a homogenous polymer model of chromatin fiber (i.e. the distribution of nucleosomes or DNA density is uniform along it). MD simulations of the homogenous polymer model are performed with and without the compartmentalization interaction to generate two ensembles containing 500 chromatin conformations of K562, denoted as the AB/homo-ensemble and the homo-ensemble.

The features of the individual contact matrices of the 5-Mb region (Chr5:109Mb-114Mb) are very similar among the four ensembles (i.e. hetero-ensemble(ATAC), AB/hetero-ensemble(ATAC), homo-ensemble, and AB/homo-ensemble), without any notable change in the boundary strength and number of domain boundaries within conformations (Fig. 2A, F, G, and Fig. S4 H to K). The similarity implies that random folding of chromatin inherently drives the extensive and random formation of insulated domains along chromatin fiber, no matter whether the nucleosome distribution along it is non-uniform or uniform. Notably, the simulations of the four ensembles do not include any information on the so-called TAD-organizing protein (i.e. cohesin), suggesting that the domain formation in single cells is likely independent of cohesin. This result is in consistent with previous FISH observation(*15*).

We next investigate the probability for each genomic position to be a domain boundary among the individual contact matrices for the four conformation ensemble. The homo-ensemble shows a nearly flat curve of the probability throughout the 5-Mb region (Fig. 2E), while the hetero-ensemble(ATAC) and AB/hetero-ensemble(ATAC) display the curves that contain evident peaks at the positions aligning with the TAD boundaries of the Hi-C matrix (Fig. 2B to D). Consequently, no discernable domain structure is observed in the ensemble-averaged contact matrices derived from the homo-ensemble (Fig. 2E), implying that the non-uniform nucleosome distribution plays a critical role in the emergence of population-averaged TADs. The ensemble-averaged contact matrix of the AB/homo-ensemble predicts some TADs boundaries, again indicating that the inclusion of the interaction for compartmentalization in the RCHC model can improve the prediction of genome organization.

Finally, our simulations on the random folding of heterogeneous and homogenous chromatin indicate that genome is randomly insulated into domains with bias introduced by the non-uniform nucleosome distribution which leads to the emergence of TADs at the population level.

### RCHC predicts the chromatin reorganization during differentiation process

To test the promise of RCHC to facilitate scientific inquiry in genome biology, we apply RCHC to predict chromatin structures of the cell types involved in the eight developmental stages of the differentiation from hematopoietic stem and progenitor cells (HSPCs) to mature immune T cells. For each stage, previous experimental study has provided the Hi-C contact-frequency matrices of the two regions enclosing the critical regulator genes of the differentiation, *Meis1* and *Bcl11b*, and the DNase-seq data of the same two regions (*27*).

We use the DNase-seq data as input to generate a conformation ensemble of each region for each stage. Ensemble-averaged contact matrices are derived from these conformation ensembles to compare to the Hi-C matrices (Fig. 5). For a quantitative comparison, we again use separation score to characterize the strength of spatial segregation at each genomic position for all the matrices in Fig. 5. The separation score profiles of the Hi-C maps reveal that the strength of the two boundaries of the TAD containing *Meis1* sharply decrease at the transition from DN3 stage to DN4, while the TAD enclosing *Bcl11b* begins to appear at DN2 stage. Previous study proposes that these abrupt changes act as a barrier to lock the cell fate into the T lineages (*27*). When we turn to the simulated ensemble-averaged contact matrices, we immediately observe that such key chromatin reorganizations is perfectly predicted by the simulated matrices. The consistence between the simulation and the Hi-C experiment result shows the promise of RCHC to investigate the chromatin reorganization involved in differentiation and cell-fate decision.

**Fig. 5.**
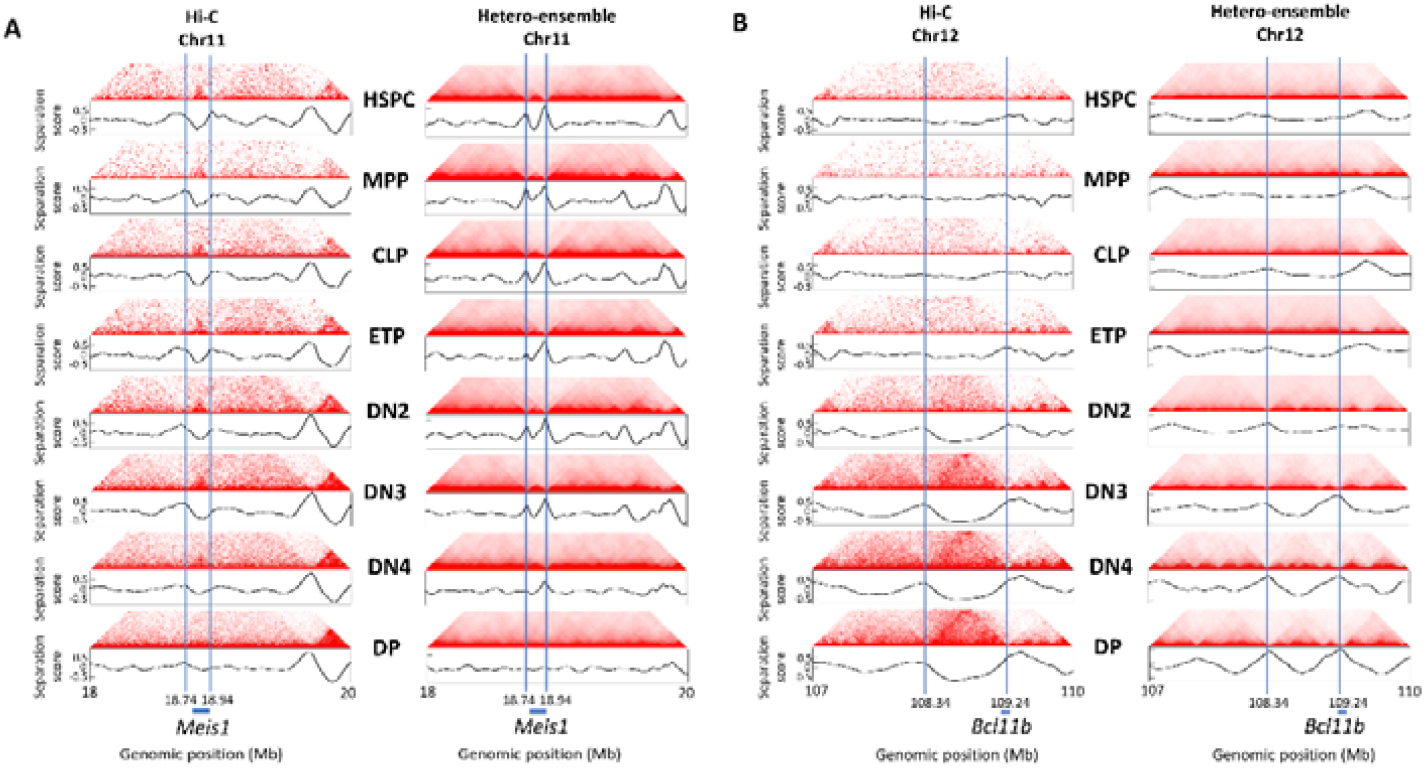
Chromatin reorganizations during the differentiation from hematopoietic stem and progenitor cells (HSPCs) to mature immune T cells. **(A** and **B)** Left: Hi-C contact-frequency matrices of the two genomic regions harboring *Meis1* (A) and *Bcl11b* (B) for eight developmental stages (HSPC, MPP, CLP, ETP, DN2, DN3, DN4, and DP) during the differentiation. Right: The corresponding simulated ensemble-average contact matrices of the two same regions. Blue lines indicate domain boundaries identified by separation score. Further details of simulated ensemble-average contact matrices are given in the supplementary materials [(*24*), sections 2].

## Discussion

We propose that the TADs emergency is underlied by random folding of chromatin fiber heterogeneous in nucleosome density. The new hypothesis contains two basic assumptions. First, since the physical process of randomly folding chromatin fiber within a nucleus inherently causes the wide and random occurrence of locally compacted regions on chromatin fiber, we assume that the random folding process itself results in the sTADs formation in single cells and the cell-to-cell variation in sTADs. Second, based on the experimental observation that chromatin regions with CTCF binding always show a non-uniform distribution of nucleosomes, we assume that non-uniform nucleosome distribution along chromatin is associated with the emergence of TADs in population-averaged Hi-C maps. To test the hypothesis, we build the RCHC model to simulate random folding of heterogeneous chromatin fiber and de novo predict chromatin structure ensemble from chromatin accessibility data (reflecting nucleosome density and from DNase-seq or ATAC-seq experiments). The validity of the RCHC model can be assessed by data from not only population-based Hi-C maps but also single-cell FISH images, demonstrating that the hypothesis behind the model indeed integrates the different features of chromatin organization revealed by the two independent experimental techniques.

The RCHC model substantially outperforms the accepted model for the TADs formation, the cohesin-CTCF mediated loop extrusion (LE) model. The inferiority of the LE model can be attributed to three fundamental factors. First, the LE model involves a priori assumption that cohesin extrudes a chromatin region into spatially insulated domain. Although the LE hypothesis has prompted a lot of experimental studies(*28-30*), no evidence has been yielded to support a cohesin-mediated extrusion process with the size of extruded region more than 100 kb. On the other hand, FISH imaging documents the widespread existence of domains in single cells depleted of cohesin, showing a clear and convincing evidence for the notion that cohesin is not required for the formation of domains in single cells(*15*). In contrast, the RCHC simulation process just follows the laws of polymer physics without any priori assumption. Second, the simulation input data of the LE model, such as the extrusion velocity and the size of an extruded region, is yet challenging for experimental study(*31, 32*), whereas the input of the RCHC model is easily available from DNase-seq and ATAC-seq experiments. Therefore, the RCHC model is more practical than the LE model for genome biology research. Third, it is reasonable to think that the LE model considers a directed process for the TADs formation based on a static and rigid genome organization, because the LE model is rooted in two facts revealed by population-based chromosome capture experiments: (i) TADs are largely cell type invariant with boundaries decorated by cohesin and CTCF; (ii) depletion of cohesin or CTCF accompanies loss of TADs and ectopic gene regulation. Such static and rigid picture might give the impression that genome organization severs as binary determinant of gene function. However, growing evidences have made it clear that both genome organization and function are random and dynamics(*7, 12, 13*). In particular, recent two experiments show that the genome organizations are similar among cell types with highly heterogeneous gene expression patterns, indicating that chromatin structure might act as a flexible foundation, rather than binary determinant of gene function(*33-35*). Our RCHC model, unlike the LE model, considers the emergence of TADs in the context of a dynamic and random genome organization. The key idea of RCHC is that genome is organized randomly with biases introduced by the DNA-encoded information such as nucleosome distribution which bring about the emergent feature of genome organization, for instance TADs.

Notably, the experimental study of genomes has progressed from inspection of single, exemplary chromatin regions to genome-wide analysis. However, most reported polymer models considering how genome is organized are limited to exemplary regions(*36*), although mechanistic modeling is regarded as useful in unveiling the mechanistic basis and functional implication of genome organization(*13*). Our RCHC model for the first time predicts mammalian chromatin organization of 10-kb resolution at genome-wide scale, with the prediction quantitatively validated by population Hi-C data and single-cell FISH data.

In addition, we build a polymer model of chromatin fiber carrying multiple DNA-encoded information including nucleosome occupancy and transcriptional activity to simulate its random folding under the self-interaction between heterochromatic regions. Such simulation simultaneously predicts domain and compartment structures. The comparison of the simulations with and without the interaction for compartment formation confirms the experimentally observed independence between local insulation and genomic compartmentalization in genome organization. Our simulations indicate that the RCHC model might server as a framework to integrate diverse DNA-encoded information for a more accurate prediction of genome organization.

Together, the RCHC model proves that genome organization is random with biases introduced by DNA-encoded information. The RCHC model might pave the way to relate 3D genome organization to multi-omics datasets and will help us better understand the relationship between genome organization and gene function.

## Author contributions

L. M. and Q. L. conceived of the study. L. M. developed the RCHC model and performed the calculations. L. M. and Q. L. performed the analysis and interpreted the results. Qiong Luo. wrote the paper with input L. M..

