## Supplementary File for "Random folding drives the emergence of topologically associating domains in chromatin three-dimensional structure"

**The PDF file includes:**

Materials and Methods

Supplementary Text

Figs. S1 to S8

References (37-38)

**1 The RCHC model**

**1.1 The polymer model of heterogeneous or homogeneous chromatin fiber**

We first partition the chromatin region of interest into *N* consecutive 10-kb length segments. Next, we model the number *k_i_* of contiguous, coarse-grained, and equal-volume beads used to represent the *i^th^* segment as a random independent Poisson variable. The possibility of *k_i_*, *P(k_i_)*, is defined as:

$P\left( k_{i} \right)=\frac{{\lambda_{i}}^{k_{i}}}{k_{i}!}e^{{-\lambda}_{i}}$	(s1)

**The definition of λ_i_:** Based on the experimental population-based chromatin accessibility data of the chromatin region of interest (i.e. DNase-seq or ATAC-seq data), we calculate the DNA accessibility signal ζ for each 10-kb segment within the chromatin region of interest. *λ_i_* is the normalized value of DNA accessibility signal of the *i^th^* 10-kb segment and is defined as:

$\lambda_{i}=\frac{\zeta_{i}}{\text{Median}(\zeta)}$	(s2)

**The determination of *k_i_*:** With the constant *λ_i_*, the value of *k_i_* is determined by using a pseudo-random number generating function capable of simulating Poisson random variables.

**The establishment of polymer model for heterogeneous chromatin fiber:** Performing the process of determining *λ_i_* and *k_i_* for each 10-kb segment in the chromatin region of interest, we obtain a certain set of *K*(*k_1_*, *k_2_*,…*k_N_*) to build a polymer model with a certain DNA density distribution for the chromatin region of interest. To model the cell-to-cell variation in the DNA density distribution (i.e. the nucleosome distribution), we generate ten distinct sets of *K*(*k_1_*, *k_2_*,…*k_N_*) to represent ten different distributions of DNA density along the same region of interest. Namely, we build ten distinct polymer models for the chromatin region of interest. Fig. S1 shows the resulted ten sets of *K*(*k_1_*, *k_2_*,…*k_N_*) for the 5-Mb chromatin regions (Chr5:109Mb-114Mb) of K562 and IMR90 as example, from population-based chromatin accessibility data.

**The establishment of polymer model for homogeneous chromatin fiber:** If the chromatin region of interest is homogeneous in DNA density, we set all the Poisson variables in *K*(*k_1_*, *k_2_*,…*k_N_*) to be 1. Namely, for homogeneous chromatin fiber, each equal-length segment is represented by one equal-volume bead.

**1.2 Molecular dynamics (MD) calculation**

Following the method mentioned in Section 1, each chromosome is represented as a polymer chain consisting of contiguous, equal-volume beads with the diameter of *b*. Moreover, each bead is same in mass with the value of *m*. The 3D chromatin structure of a chromatin region of interest is simulated by numerical integration starting from an initially random conformation with the step size $\Delta t$ defined as:

$\Delta t=\sqrt{0.2b*\frac{F_{max}}{m}}$	(s3)

where $F_{max}$ is the maximum force among the forces applied on beads of the polymer model for the represented chromatin region of interest. The diameter *b* and the mass *m* of each bead are set to 1. At each step during the simulation, the velocity of each bead is randomly generated following the Boltzmann distribution.

**Total potential energy in 3D conformation, excluding the interaction for the compartments formation:** We define the total potential energy with three terms that are described below:

(1) Harmonic oscillator potential energy, $E_{har}\left( r_{n,n+1} \right)$, is introduced to ensure the connectivity of chromosome backbone and is calculated according to the formula as the following:

$E_{har}\left( r_{n,n+1} \right)={\frac{1}{2}k_{b}\left( r_{n,n+1}-b \right)}^{2}$	(s4)

where $r_{n,n+1}$ is the Euclidean distance between pair of the sequentially adjacent *n^th^* and (*n*+1)*^th^* beads. The equilibrium distance between the *n^th^* and (*n*+1)*^th^* beads is approximately equal to the bead diameter *b*. The force constant $k_{b}$ is defined as:

$k_{b}={\frac{200kT}{b^{2}}}$	(s5)

where $k$ is the Boltzmann constant, *T* the temperature, and *b* the diameter of bead with the value of 1. Here, *kT* is considered as a unit amount of energy.

(2) Repulsive potential energy, $E_{WCA}\left( r_{n,m} \right)$, is introduced to avoid spatial overlapping between pair of the non-sequentially-adjacent *n^th^* and *m^th^* beads and is calculated according to the following:

$E_{WCA}\left( r_{n,m} \right)=\left\{ \begin{matrix} 4\varepsilon\left( \left( \frac{\sigma}{r_{n,m}} \right)^{12}-\left( \frac{\sigma}{r_{n,m}} \right)^{6} \right), if r_{n,m}\leq2^{1/6}\sigma\\ -\varepsilon, else \end{matrix} \right.$	(s6)

Where $r_{n,m}$ represents the Euclidean distance between the *n^th^* and the *m^th^* beads and $\sigma$ and $\varepsilon$ are the two constants with $\sigma=b$ and $\varepsilon=\frac{1}{2}kT$.

(3) Sphere restraint potential energy,$E_{sphere}\left( r_{n} \right)$, is introduced to constrain the folding of the polymer model of chromatin in a confined sphere space like cell nucleus. During the structure calculation, the center of the sphere space is fixed as origin of coordinates (0,0,0). The radius of the sphere $r_{s}$ is defined as:

$r_{s}=\frac{1}{2}b\left( N/0.35 \right)^{1/3}$	(s7)

where *N* is the total number of beads in the polymer model and *b* the diameter of each bead with the value of 1.

Based on the definition of the sphere space, sphere restraint potential energy,$E_{sphere}\left( r_{n} \right)$, is calculated according to the formula:

$E_{sphere}\left( r_{n} \right)=\left\{ \begin{matrix} \frac{r_{n}-r_{s}}{b}kT, if r_{n}\geq r_{s} \\ 0, else \end{matrix} \right.$	(s7)

Where $r_{n}$ is the Euclidean distance between the *n^th^* bead and the center of the sphere space (i.e. origin of coordinates).

**Potential energy to describe the interaction for the A/B compartments formation:** Previous studies have shown that self-interaction between heterochromatic regions leads to compartmentalization of chromatin into A/B compartments. Therefore, the attraction potential energy, $E_{B}\left( r_{n,m} \right)$, is introduced to describe the attractive interaction between pair of the *n^th^* and *m^th^* beads belonging to transcriptionally inactive regions and is defined as:

$E_{B}\left( r_{n,m} \right)=\left\{ \begin{matrix} 10\varepsilon\left( \ln\left( \frac{r_{n,m}}{b} \right)-\frac{r_{n,m}}{2b} \right), if r_{i,j}\leq2b \\ 10\varepsilon(ln2-1), else \end{matrix} \right.$	(s8)

where $r_{n,m}$ represents the Euclidean distance between the *n^th^* and *m^th^* beads, *b* the diameter of each bead with the value of 1, and $\varepsilon$ the constant equal to 1/2$kT$. Notably, the method for identifying beads belonging to transcriptionally inactive regions is shown in Section 6.

**1.3 Hierarchical calculation strategy**

As mentioned in section 1.1, we equally partition the chromatin region of interest into *N* consecutive 10-kb segments and build a bead-on-a-string representation of the region. We define the resolution of the representation as the sequence length of individual partitioned segments, namely 10 kb. To rapidly achieve the conformation of the chromatin region of interest at 10-kb resolution, we employ a hierarchical protocol which includes the following steps:

(1) We first determine the number *k* of beads used to represent each 10-kb segment according to the method in Section 1 and obtain a certain *K* (*k_1_*, *k_2_*,…*k_N_*) to build a bead-on-a-string representation for the chromatin region of interest at 10-kb resolution. We next sequentially merge two adjacent 10-kb segments into one 20-kb segment. Based on the numbers of beads (*k_i_* and *k_i+1_*) of the two *i^th^* and (*i+1*)*^th^* merged parent 10-kb segments, we define the number of beads used to represent the resulted 20-kb segments, denoted as $\bar{k_{i\_20}}$ , according the following:

$\bar{k_{i\_20}}=\frac{k_{i}+k_{i+1}}{2}$	(s9)

After the determination of bead numbers for all resulted 20-kb segments, we obtain the bead-on-a-string representation for the chromatin region of interest at 20-kb resolution. In the same manner, we generate the representation for the same region of 40-kb resolution from the representation of 20-kb resolution. We continue the procedure to generate the representation at the lower resolution, including 80-kb, 160-kb, 320-kb, 640-kb, 1280-kb, 2560-kb, and 5120-kb resolutions. The aim of this step is to determine the number of beads used to represent the chromatin region of interest at lower resolutions based on the given *K* (*k_1_*, *k_2_*,…*k_N_*).

(2) Initial 3D structure of the representation at 5120-kb resolution is randomly generated using self-avoiding random walk and the numerical integration following the procedure described Section 2 is performed for 200,000 steps to generate the simulated 5120-kb conformation. Based on the coordinates of beads in the simulated 5120-kb conformation, initial coordinates of beads in the representation of 2560-kb resolution can be determined by using linear interpolation. Namely, the initial conformation of 2560-kb resolution for the structure calculation is derived from the output of the structure calculation at 5120-kb resolution. In the same manner, the structure calculations are performed at 2560-kb, 1280-kb, 640-kb, 320-kb, 160-kb, 80-kb, 40-kb, 20-kb and finally 10-kb resolutions with the initial structure of each stage generated from the previous round of structure calculations. Such hierarchical simulation is performed for 50 times from 50 different initial 3D structures at 5120-kb resolution to finally generate 50 different simulated conformations of the chromatin region of interest at 10-kb resolution for a given *K* (*k_1_*, *k_2_*,…*k_N_*).

**2 Conformation ensembles involved in the study**

In this study, we generate six distinct conformation ensembles for two human cell lines: K562 erythroleukemia and IMR90 lung fibroblasts. Moreover, we generate conformation ensembles for the eight cell types involved in the differentiation from hematopoietic stem and progenitor cells (HSPCs) to mature immune T cells.

**Six conformation ensembles for K562 cell type:** The chromatin region of interest includes 22 autosomes and an X-chromosome of K562 human erythroleukemia cell. The six simulated conformation ensemble are:

(1) Hetero-ensemble(ATAC) and Hetero-ensemble(DNase): We equally partition the region of interest into consecutive 10-kb segments and build ten distinct polymer model of the region according the method described in Section 1.1, with the DNA accessibility data from the Gene expression Omnibus (GEO). Specifically, the ATAC-seq data (GEO:GSE170214) is used to generate Hetero-ensemble(ATAC) and the DNase-seq data (GEO:GSM1008601, GSM1008580, GSM1008567, GSM816655, GSM1008602, and GSM1008558) is used to generate Hetero-ensemble(DNase).

For each bead-on-a-string polymer model, we perform the molecular dynamics (MD) calculation with the total potential energy including three terms (i.e. Harmonic oscillator potential energy, Repulsive potential energy, and Sphere restraint potential energy, described in Section 1.2 and 1.3) to simulate its random folding in a confined space for 50 times starting from 50 different random initial structures. Following the same process, we respectively produce two ensembles consisting of 500 conformations of the chromatin region of interest at the resolution of 10 kb, for both ATAC-seq and DNase-seq input data. We denote the two ensembles as hetero-ensemble(ATAC) and hetero-ensemble(DNase), because the ensembles are derived from random folding of chromatin fiber heterogeneous in terms of DNA density.

(2) AB/hetero-ensemble(ATAC) and AB/hetero-ensemble(DNase): The procedure for generating the AB/hetero-ensemble(ATAC) and AB/homo-ensemble(DNase) is very similar to that for Hetero-ensemble(ATAC) and Hetero-ensemble(DNase), except for the difference in the definition of the total potential energy of a 3D conformation in the MD calculation. For convenience, we collectively refer to hetero-ensemble(ATAC) and hetero-ensemble(DNase) as hetero-ensembles and collectively refer to AB/hetero-ensemble(ATAC) and AB/hetero-ensemble(DNase) as AB/hetero-ensembles. Comparing to the hetero-ensembles, the potential energy associated with the self-interaction between heterochromatic regions is added into the total potential energy of a 3D conformation in the AB/hetero-ensembles.

(3) Homo-ensemble and AB/home-ensemble: By setting all the Poisson variables *k_i_* in *K* (*k_1_*, *k_2_*,…*k_N_*) equal to 1, we build a homogenous polymer model consisting of same beads. For the homogenous polymer model, the MD calculation with the total potential energy including three terms (i.e. Harmonic oscillator potential energy, Repulsive potential energy, and Sphere restraint potential energy) is performed for 500 times starting from 500 different random initial structures and generate a conformation ensemble, denoted as homo-ensemble. Similarly, by adding the self-interaction between heterochromatic regions into the total potential energy of a 3D conformation, the same procedure for generating the homo-ensemble produces the AB/homo-ensemble.

Notably, the analysis of hetero-ensemble(DNase), AB/hetero-ensemble(DNase), homo-ensemble and AB/home-ensemble are shown in Fig. S4 and S5.

**Conformation ensembles for IMR90 cell type:** To investigate the question whether the RCHC model is applicable to other cell types except for K562 erythroleukemia cell type, we generate similar conformation ensembles for IMR90 cell type with the ATAC-seq data (GEO:GSE169767) as input. The analysis of the ensembles is displayed in Fig. S6 to S8.

**Conformation ensembles for cell types involved in the eight developmental stages of the differentiation from HSPCs to mature immune T cells:** There are two chromatin region of interest. One is the 10-Mb genomic region (Chr11:15Mb-25Mb) containing the regulator gene *Meis1* and the other is the 10-Mb genomic region (Chr12:105Mb-115Mb) enclosing the regulator gene *Bcl11b*. The eight cell types involves the developmental stages of the differentiation are hematopoietic stem and progenitor cells (HSPCs), multipotent progenitor (MPP), common lymphoid progenitor (CLP), early T precursor(ETP), CD4 and CD8 double-negative 2 (DN2), DN3, DN4, and double-positive (DP) cells, respectively. For each region at each stage, we apply the RCHC model (the total potential energy includes three terms, i.e. Harmonic oscillator potential energy, Repulsive potential energy, and Sphere restraint potential energy) to generate an ensemble containing 500 conformations at 10-kb resolution, with the DNase-seq experimental data (GEO:GSE79422) as input. Therefore, we generate 16 ensembles.

**3 Contact matrix**

**Individual contact matrix:** All the ensembles mentioned in Section 2 contain 500 individual 3D conformations of 10-kb resolution. As described in Section 1, the chromatin fibers of each conformation of 10-kb resolution are divided into 10-kb segments with each segment represented by a few of equal-volume beads. For each conformation, we measure the distance between the centroids of each pair of 10-kb segments. Based on the measurement, we build a contact matrix ***C*** for each conformation at the resolution of 10-kb, in which matrix element *c_ij_* is equal to 1 if the distance between the *i^th^* and *j^th^* segments is less than 4 bead diameters and *c_ij_* is zero when the distance is larger than 4 bead diameters.

**Ensemble-averaged contact matrix:** For each ensemble, we average the individual contact matrices across 500 conformations and obtain the ensemble-averaged contact matrix ***C***^*^ which is defined as:

$c_{ij}^{*}=\frac{A_{ij}}{\sqrt{A_{i}*A_{j}}}$	(s10)

where $A_{ij}$ is the element of the matrix ***A*** which is generated by directly average individual contact matrices ***C*** across 500 conformations. The matrix ***A*** is defined as:

$A_{ij}=\frac{1}{500}\sum_{n=1}^{500} c_{ij,n}$	(s11)

Moreover, the $A_{i}$ and $A_{j}$ in the equation (s10) are calculated according to the formula below:

$A_{i}=\sum_{j} A_{ij}$ and $A_{j}=\sum_{i} A_{ij}$	(S12)

The ensemble-averaged contact matrix ***C***^*^ can be compared to the population-averaged Hi-C contact matrix.

**4 Distance matrix**

**4.1 Distance matrices calculated from the hetero-ensembles and AB/hetero-ensembles**

**Individual distance matrix:** The calculation of the individual distance matrix at 30-kb resolution from individual conformations of 10-kb resolution is performed through two steps:

(1) Since each conformation in the ensembles mentioned in Section 2 is consisting of consecutive 10-kb segments with each segment represented by a few of equal-volume beads, we sequentially merge three adjacent 10-kb segments into one 30-kb segments.

(2) We measure the distance between the centroids of each pair of 30-kb segments and build an individual distance matrix for each conformation.

**Ensemble-averaged distance matrix:** For each ensemble, we directly average the individual distance matrices across 500 conformations and obtain the ensemble-averaged distance matrix.

**4.2 Distance matrix calculated from the FISH data**

Bintu *et al.* used Multiplexed super-resolution fluorescence in situ hybridization (FISH) to image 65 segments in the 2-Mb chromatin region (Chr21:29.38Mb-31.33 Mb) of K562 (*15*). A total number of 13,997 sample cells are imaged. Based on the FISH data, the average distance of each segment pair can be calculated over all the imaged cells and an averaged distance matrix can be obtained (shown in Fig. 3E).

**5 Domain boundaries in contact matrix**

**5.1 Separation score for identification of domain boundaries**

In our study, we use separation score to identify domain boundaries in all contact matrices as well as population-averaged Hi-C contact matrices. Here, we use population-averaged Hi-C contact matrix ***C*** as an example to illustrate the identification of domain boundaries.

First, we transform the populated-averaged Hi-C contact matrix ***C*** into an arrowhead matrix ***M*** which is defined as:

$M_{i,i+d}=\frac{C_{i,i-d}-C_{i,i+d}}{C_{i,i-d}+C_{i,i+d}}$	(s13)

As shown in Fig. S2, this transformation replaces domains with an arrowhead-shaped motif.

Second, we define separation score which quantify the degree of spatial separation between upstream and downstream chromatin of a position based on the arrowhead matrix ***M***. As shown in Fig. S2B, we construct two edge-shared congruent right triangles, denoted as A and B, at each position along the diagonal line of the matrix ***M***. Separation score of the *i^th^* position, denoted as *S_i_*, is defined according to the following:

$S_{i}=\frac{1}{N_{A}}\sum_{i,j\in A} M_{ij}-\frac{1}{N_{B}}\sum_{i,j\in B} M_{ij}$	(s14)

where *N*_A_ and *N*_B_ are the total number of matrix elements in the triangles A and B, respectively. In our calculation, the length of horizontal edge of the two congruent right triangles is 250-kb and the length of the vertical edge (or shared edge) is 500-kb.

Finally, we draw the curve of separation score throughout the chromatin region of interest. The positions that are identified as domain boundaries should stratify two criteria. First, the boundary position should correspond to a score peak and simultaneously show higher score value than any other positions within the 100 kb regions on either side of the position. Second, the separation score of the boundary should be higher than the average score value throughout the region.

**5.2 Identification of pair of overlapping boundaries among ensemble-average contact matrices and Hi-C contact matrix**

To assess the performance of the RCHC model, we compare the genomic positions of domain boundaries among Hi-C contact frequency matrix and the ensemble-average contact matrices derived from the hetero-ensembles and AB/hetero-ensembles of K562. Similar comparisons are also carried out for IMR90. We regard two boundaries from different contact matrices as a pair of overlapping boundaries, if the distance between their genomic positions is less than 100 kb.

**6 Identification of A/B compartments**

The identification of compartment structures from population-averaged Hi-C contact matrix, ensemble-averaged contact matrices of the AB/hetero-ensembles and hetero-ensembles are calculated following a similar algorithm described in the previous work(*37*). We present the steps for the identification of compartments from the Hi-C contact-frequency matrix of 10-kb resolution, as an example.

(1) We first transform the resolution of the parent Hi-C matrix from 10 kb to 200 kb. Next, we normalize the 200-kb Hi-C contact frequency matrix through dividing each matrix element by the genome-wide average contact frequency for bin pairs at the same genomic distance and obtain the normalized Hi-C matrix (Fig. S3).

(2) We calculate the correlation matrix ***P***, in which the element *P_ij_* describes the Pearson correlation between the *i*^th^ and *j*^th^ rows of the normalized Hi-C matrix. The correlation matrix shows many large blocks of enriched and depleted interactions, like plaid patterns (Fig. S3).

(3) Based on the correlation matrix ***P***, each chromosome is partitioned into two types of regions using spectral clustering algorithm(*38*). Between these two types of regions, the one with higher overlap with the H3K4me3 enriched regions is defined as compartment A and the other one is defined as compartment B.

**6 Data source**

The experimental data used in our analysis was taken from previously published work, as elaborated below:

| Data type | Accession number | Reference |
| --- | --- | --- |
| Populated Hi-C data for IMR90 and K562 | GEO:GSE63525 | S. S. P. Rao *et al.*, A 3D Map of the Human Genome at Kilobase Resolution Reveals Principles of Chromatin Looping (vol 159, pg 1665, 2014). *Cell* **162**, 687-688 (2015). |
| ATAC-seq data for IMR90 | GEO:GSE169767 | I. Dunham *et al.*, An integrated encyclopedia of DNA elements in the human genome. *Nature* **489**, 57-74 (2012). |
| H3K4me3 ChIP-seq for IMR90 | GEO:GSM469970,  GEO:GSM521901 |  |
| H3K4me3 ChIP-seq for K562 | GEO:GSM733680 |  |
| ATAC-seq data for K562 | GEO:GSE170214 | I. Dunham *et al.*, An integrated encyclopedia of DNA elements in the human genome. *Nature* **489**, 57-74 (2012). |
| DNase-I hypersensitivity sites for K562 | GEO:GSM1008601, GEO:GSM1008580,  GEO:GSM1008567,  GEO:GSM816655,  GEO:GSM1008602,  GEO:GSM1008558 |  |
| 3D FISH data for K562 and IMR90 |  | B. Bintu *et al.*, Super-resolution chromatin tracing reveals domains and cooperative interactions in single cells. *Science* **362**, 419-+ (2018). |
| Hi-C data and DNase-I hypersensitivity sites data for eight developmental stages from HPSC cells | GEO:GSE79422 | G. Q. Hu *et al.*, Transformation of Accessible Chromatin and 3D Nucleome Underlies Lineage Commitment of Early T Cells. *Immunity* **48**, 227-+ (2018). |

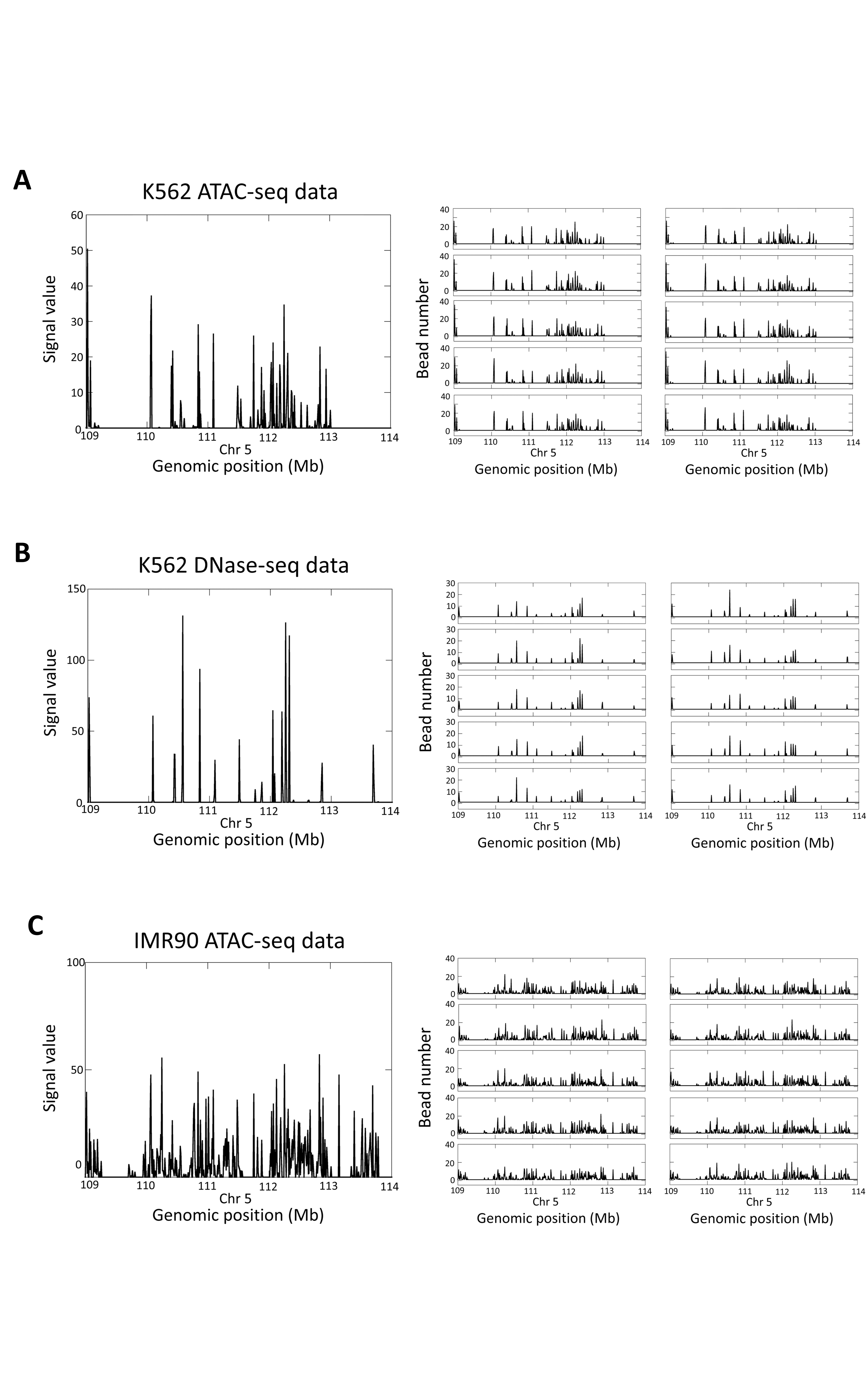

**Fig. S1. Different sets of random independent Poisson variables, *K*(*k_1_*, *k_2_*,…*k_N_*), resulted from population-based chromatin accessibility data.** Left: the chromatin accessibility data from ATAC-seq or DNase-seq experiments; right: the resulted ten sets of Poisson variables. (**A** and **B**) for K562 with experimental ATAC-seq (A) and DNase-seq (B) data as input, respectively; (**C**) for IMR90 with ATAC-seq data as input. Sources of the experimental data are listed in the *Data source* section.

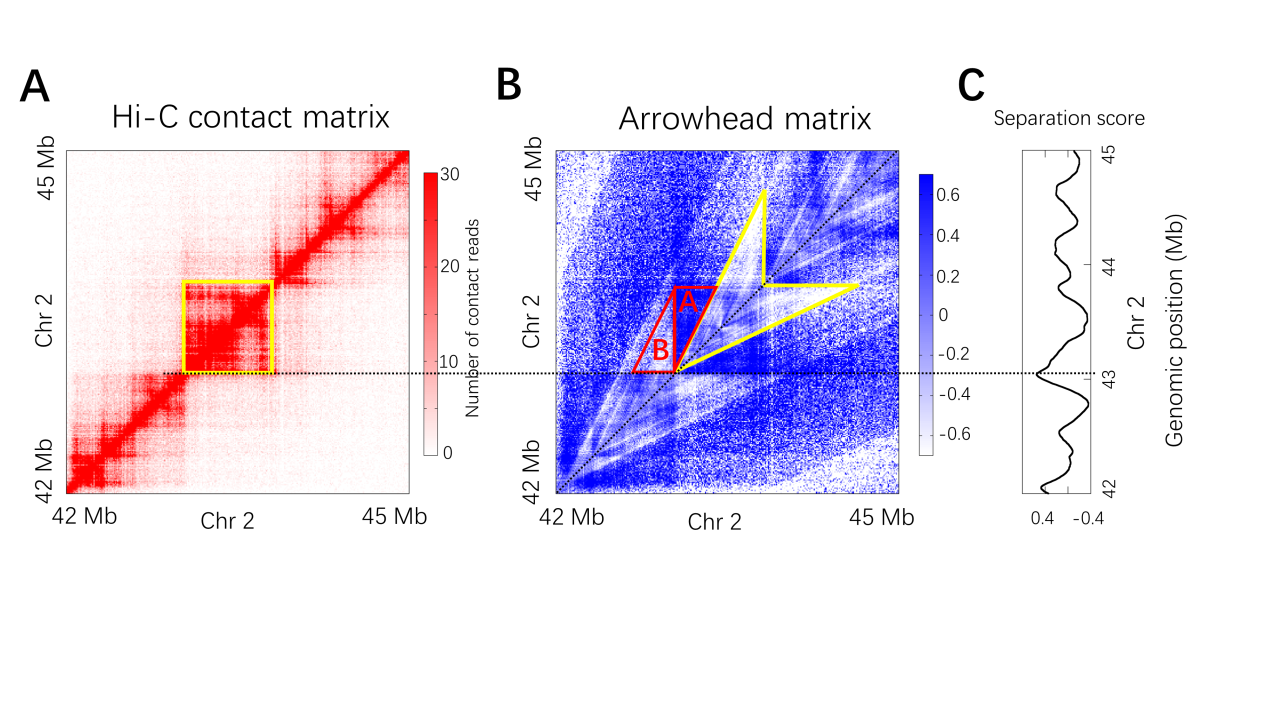

**Fig. S2.** **A scheme of the definition of separation score.** (**A**) Population-based Hi-C contact frequency matrix for the 3-Mb region (Chr2:42Mb-45Mb) of K562 at 10-kb resolution. (**B**) The arrowhead matrix *M* where the arrowhead-shaped motif highlighted by yellow corresponds to the yellow highlighted domain square in (A). (**C**) Separation score of a position along the diagonal of the arrowhead matrix is defined based on the two constructed edge-shared congruent right triangles at that position. As an example, such two edge-shared congruent right triangles of the position corresponding to a domain boundary are highlighted by red in (B). The length of horizontal edge of the two congruent right triangles is 250-kb and the length of the vertical edge (or shared edge) is 500-kb.

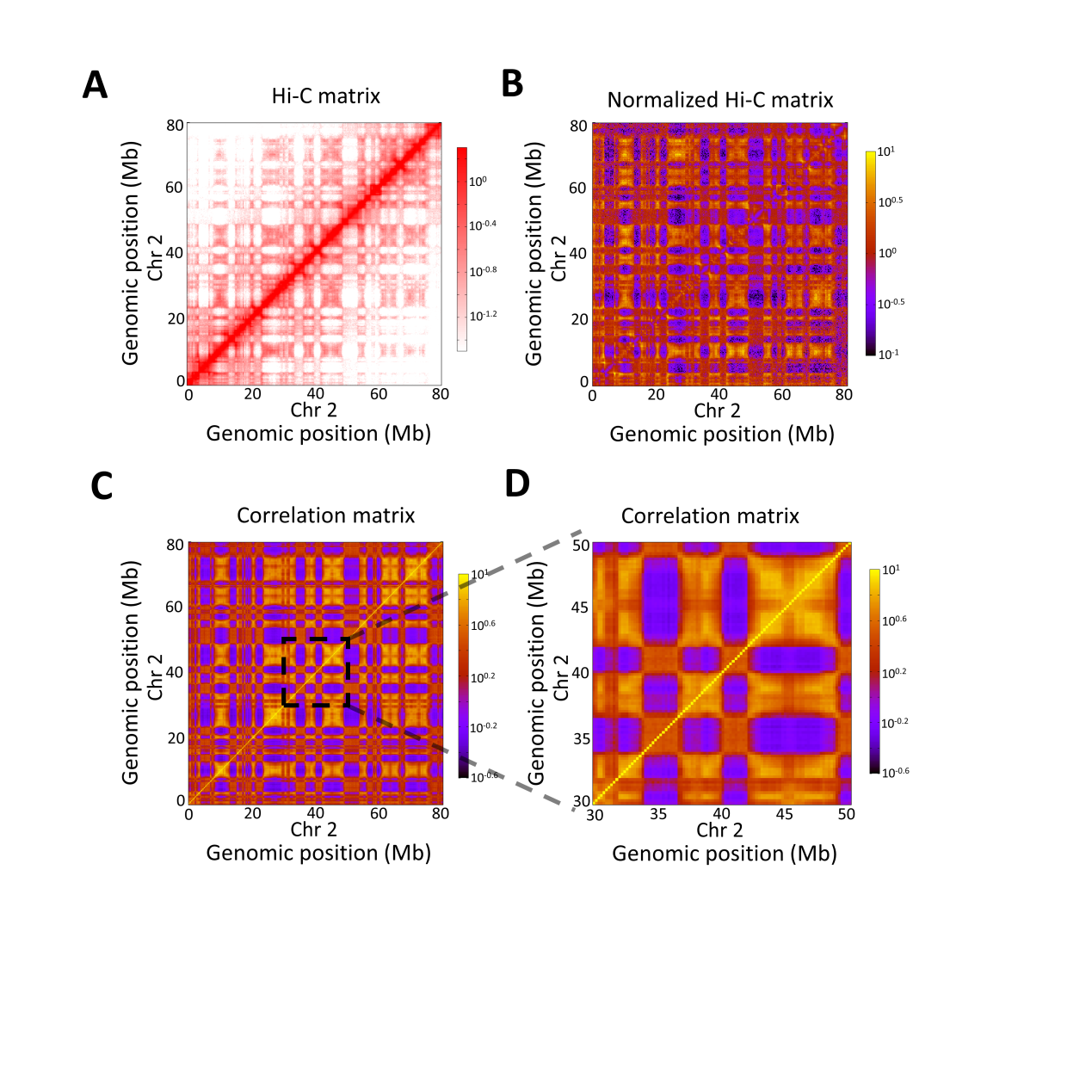

**Fig. S3. Identification of compartments from the Hi-C contact-frequency matrix.** (**A**) Hi-C matrix of the region (Chr2:0Mb -80Mb) of K562 at 200-kb resolution. (**B**) Normalized Hi-C contact frequency matrix at 200-kb resolution, calculated through dividing each matrix element in (A) by the genome-wide average contact frequency for bin pairs at the same genomic distance. (**C**) Correlation matrix calculated from (B) illustrates large blocks of enriched and depleted interactions. The plaid pattern indicates the presence of two compartments. (**D**) An expanded view of the region (Chr2: 30Mb -50Mb).

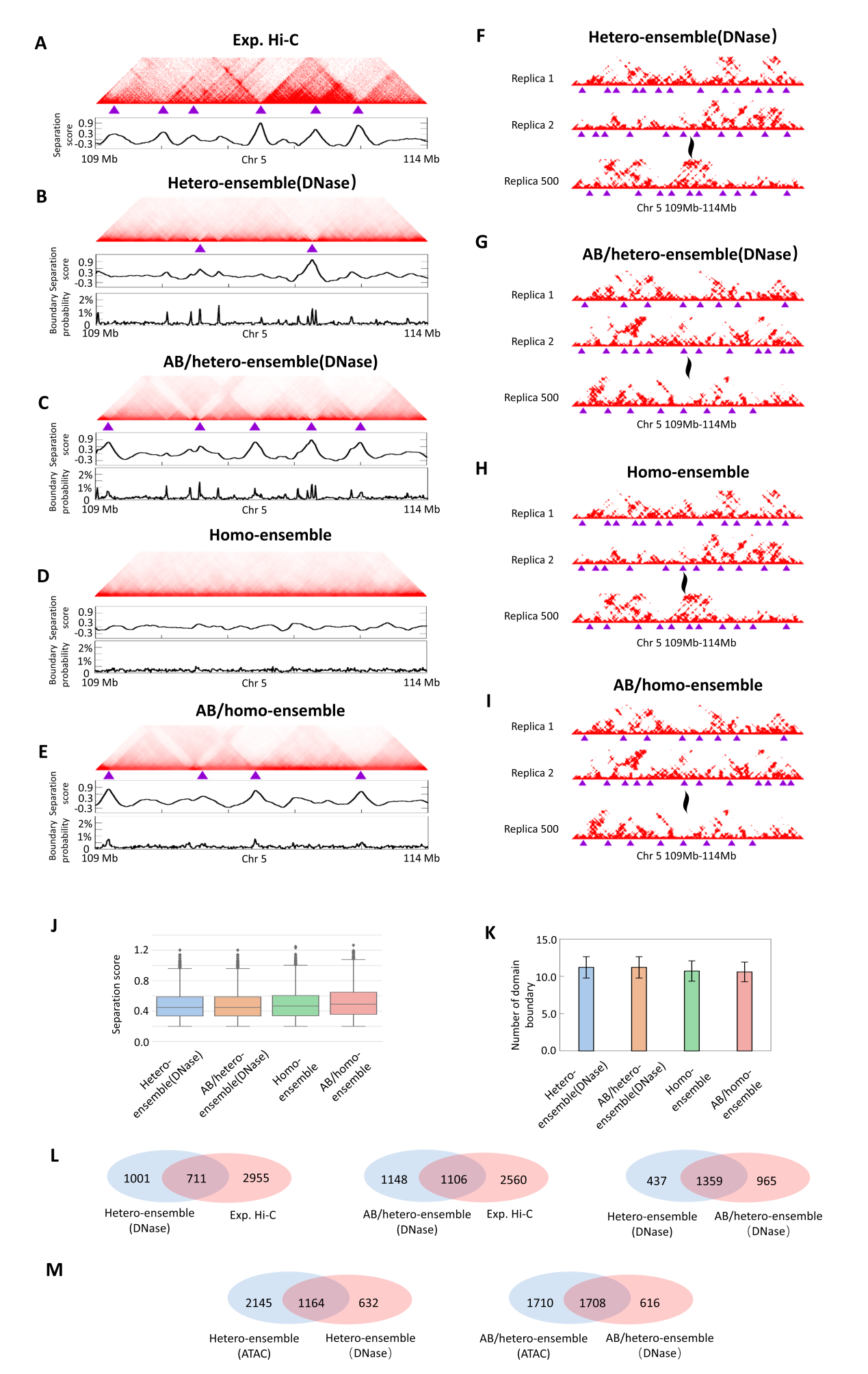

**Fig. S4. Chromatin structure ensembles of K562 from DNase-seq data**. (**A**) Top: Population-averaged Hi-C contact matrix for the 5-Mb chromatin region at 10-kb resolution. Bottom: Separation sore for each genomic position in the Hi-C matrix. (**B** to **E**) Top: The 10-kb resolution ensemble-averaged contact matrices of the 5-Mb chromatin region for (B) hetero-ensemble(DNase) describing random folding of chromatin fiber heterogeneous in DNA density using DNase-seq data as input;(C) AB/hetero-ensemble(DNase) describing the same random folding process as hetero-ensemble(DNase) but under the interaction for A/B compartment formation; (D) homo-ensemble describing random folding of homogeneous chromatin fiber; (E) AB/homo-ensemble describing random folding of homogeneous chromatin fiber under the interaction for A/B compartment formation. Middle: Separation sore for each genomic position in the ensemble-averaged contact matrix. Bottom: Probability (fraction of the 500 individual conformations) for each genomic position to appear as a domain boundary. Purple triangles denote positions of domain boundaries identified by separation score. (**F to I**) Individual contact matrices of the 5-Mb chromatin region (Chr5:109Mb-114 Mb) in K562 at 10-kb resolution, calculated from individual conformations within the four simulated ensembles: hetero-ensemble(DNase) (F); AB/hetero-ensemble(DNase) (G); homo-ensemble (H); AB/homo-ensemble (I). Purple triangles denote positions of domain boundaries identified by separation score. (**J**) Boxplots showing separation scores of domain boundaries in the individual contact matrices of the 5-Mb region. (**K**) The number of domain boundaries identified from each individual contact matrix of the 5-Mb region. (**L**) Overlap in domain boundaries throughout all autosomes and X-chromosome of K562 among the population-averaged Hi-C contact matrix, the ensemble-averaged contact matrices of hetero-ensemble(DNase) and AB/hetero-ensemble (DNase). (**M**) Overlap in domain boundaries throughout all autosomes and X-chromosome of K562 among the ensemble-averaged contact matrices between hetero-ensemble(ATAC) and hetero-ensemble(DNase) (left); AB/hetero-ensemble (ATAC) and AB/hetero-ensemble (DNase) (right). Sources of the Hi-C data are listed in the *Data source* section.

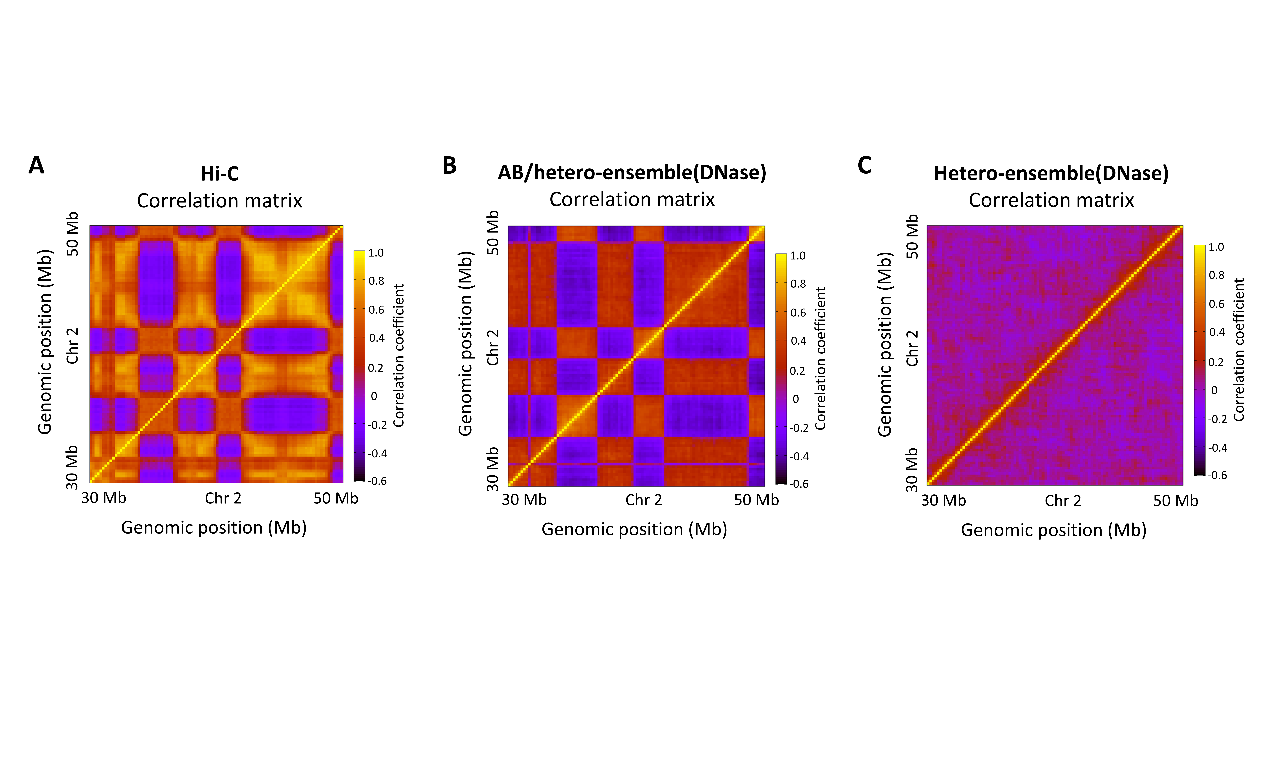

**Fig. S5. Prediction of compartment structures for K562 using DNase-seq data.** (**A**) Pearson correlation matrix of the 20-Mb region (Chr2:30Mb-50 Mb) of K562 derived from the Hi-C contact-frequency map. The plaid patterns are signatures of compartments. (**B** and **C**) Pearson correlation matrices derived from ensemble-averaged contact matrices of the AB/Hetero-ensemble(DNase) (B) and Hetero-ensemble(DNase) (C). The simulation on the random folding of heterogeneous chromatin fiber with the inclusion of the interaction between heterochromatic regions indeed quantitatively predicts compartment structures. Sources of the Hi-C data are listed in the Data source section.

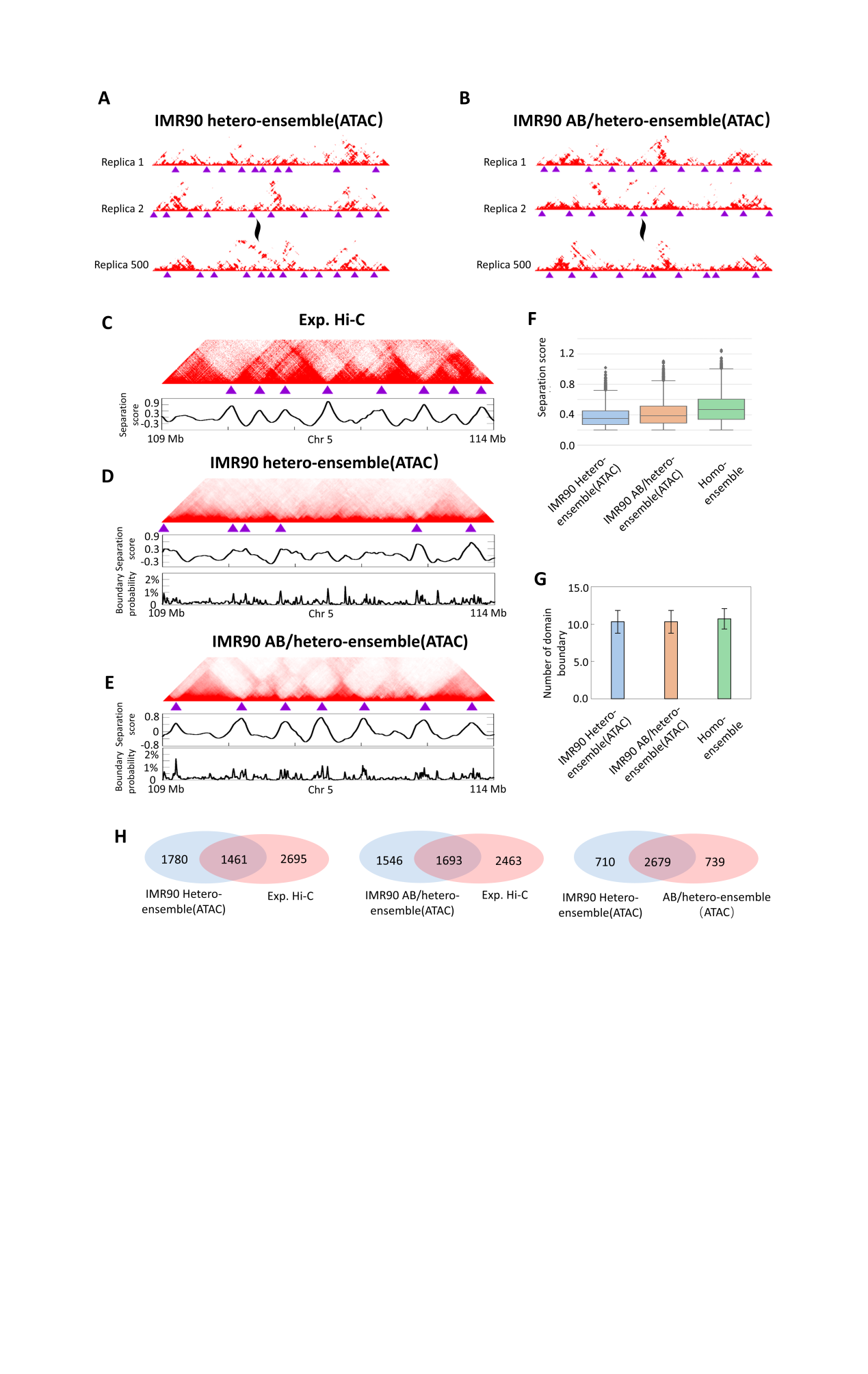

**Fig. S6. Chromatin structure ensembles of IMR90 from ATAC-seq data**. (**A** and **B**) Individual contact matrices of the 5-Mb chromatin region (Chr5:109Mb-114Mb) in IMR90 at 10-kb resolution, calculated from individual conformations within the two simulated ensembles: IMR90 hetero-ensemble(ATAC) (A) and IMR90 AB/hetero-ensemble(ATAC) (B). (**C**) Top: Population-averaged Hi-C contact matrix for the 5-Mb chromatin region at 10-kb resolution. Bottom: Separation sore for each genomic position in the Hi-C matrix. (**D** and **E**) Top: The 10-kb resolution ensemble-averaged contact matrices of the 5-Mb chromatin region for IMR90 hetero-ensemble(ATAC) (D) and IMR90 AB/hetero-ensemble(ATAC) (E). Purple triangles in (A to E) denote positions of domain boundaries identified by separation score. Middle: Separation sore for each genomic position in the ensemble-averaged contact matrix. Bottom: Probability (fraction of the 500 individual conformations) for each genomic position to appear as a domain boundary. (**F**) Boxplots showing separation scores of domain boundaries in the individual contact matrices of the 5-Mb region. (**G**) The number of domain boundaries identified from each individual contact matrix of the 5-Mb region. (**H**) Overlap in domain boundaries throughout 22 autosomes and X-chromosome of IMR90 among the population-averaged Hi-C contact matrix, the ensemble-averaged contact matrices of IMR90 Hetero-ensemble(ATAC) and IMR90 AB/hetero-ensemble(ATAC). Sources of the Hi-C data are listed in the Data source section.

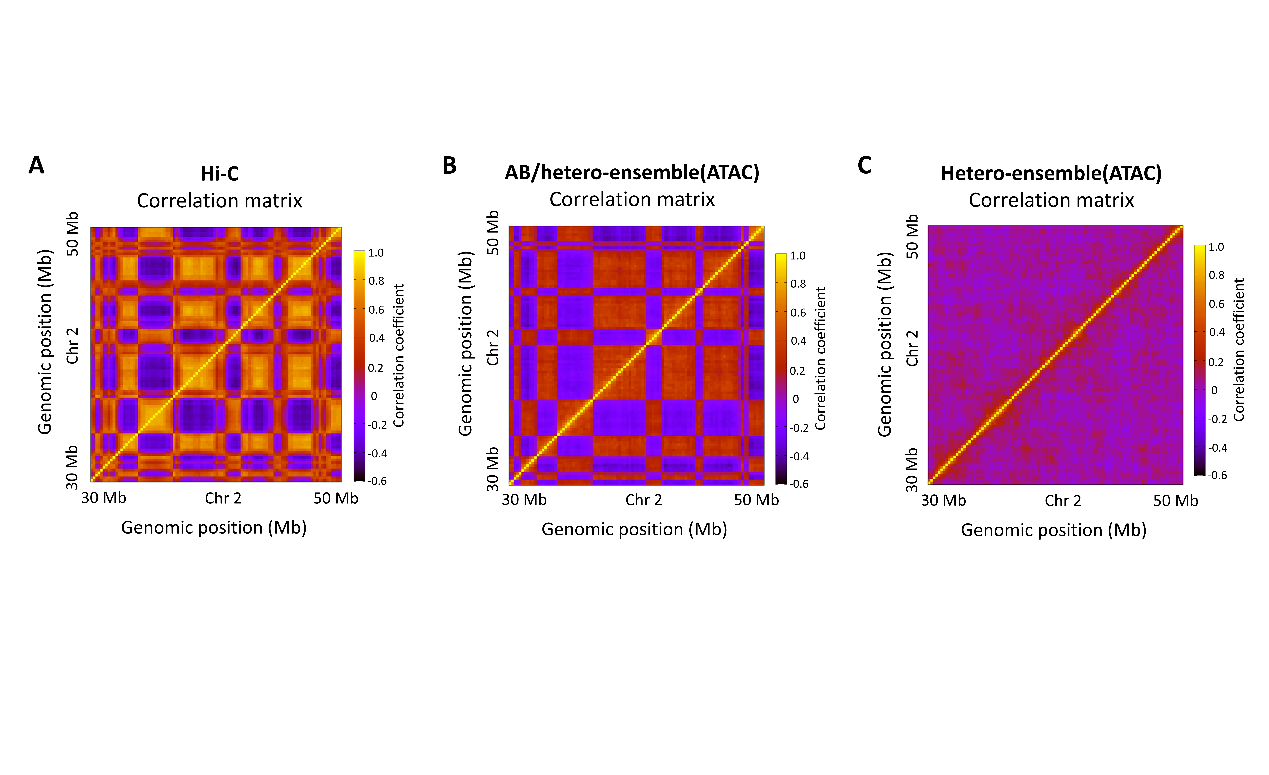

**Fig. S7. Prediction of compartment structures for IMR90 using ATAC-seq data.** (**A**) Pearson correlation matrix of the 20-Mb region (Chr2:30Mb-50Mb) of IMR90 derived from the Hi-C contact-frequency map. The plaid patterns are signatures of compartments. (**B** and **C**) Pearson correlation matrices derived from ensemble-averaged contact matrices of the AB/Hetero-ensemble(ATAC) (B) and Hetero-ensemble(ATAC) (C). Sources of the Hi-C data are listed in the Data source section.

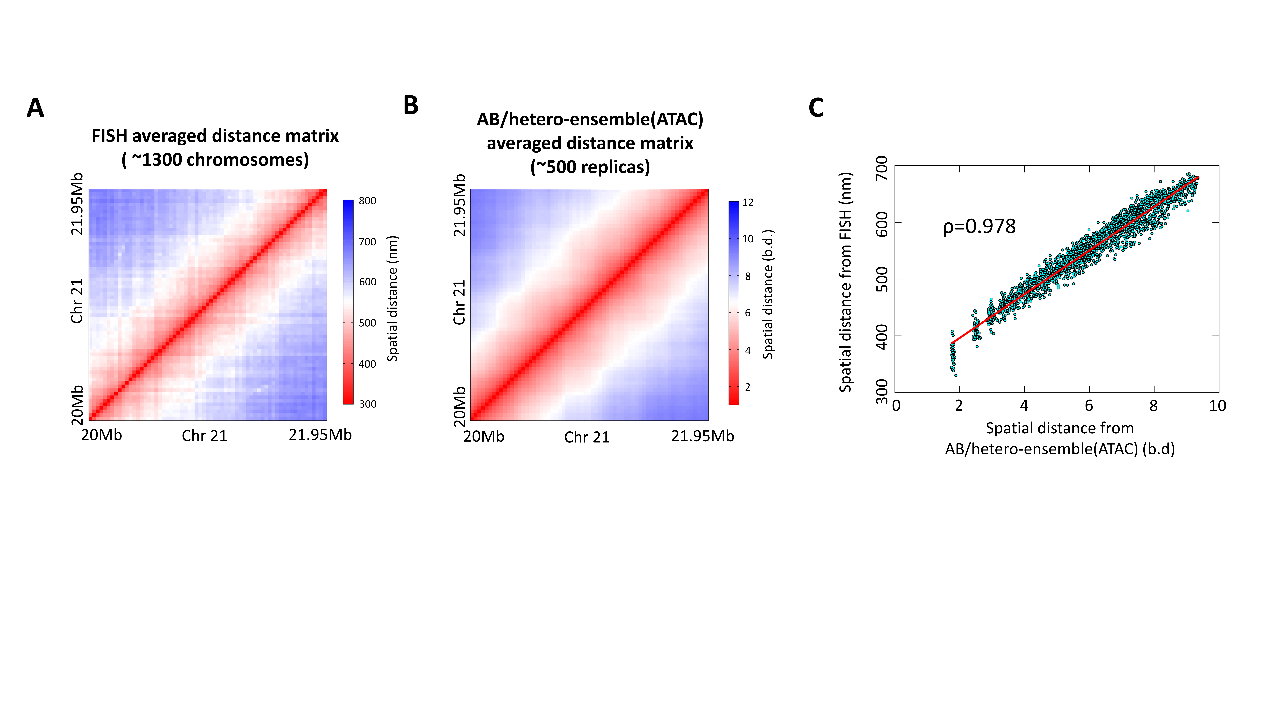

**Fig. S8. Validation of chromatin structure ensembles of IMR90 with FISH data.** (**A** to **D**) Left: the experimental averaged distance matrix at 30-kb resolution across FISH images of the same region. The 30-kb resolution averaged distance matrices of the 2-Mb region of IMR90 for experimental FISH data (A) and AB/hetero-ensemble(ATAC) (B). (**C**) Correlation between the distance matrices shown in (A) and (B).
